# Unifying phylogenetic traversal and deep learning to guide tree exploration

**DOI:** 10.64898/2026.01.14.699358

**Authors:** Lena Collienne, Harry Richman, David H. Rich, Mary Barker, Chris Jennings-Shaffer, Frederick A. Matsen

## Abstract

Deep learning offers hope for more efficient phylogenetic inference methods. However, it has yet to have the transformative effect on phylogenetics that it has had in other fields. Here we present a novel approach that combines deep learning with concepts behind current successful phylogenetic algorithms. Specifically, we give the deep learning algorithm access to the output of a phylogenetic dynamic program on the sequence alignment, rather than the raw sequence alignment. The algorithm then learns features based on these phylogenetically processed versions of the sequence data, providing information to guide local tree search. For this paper, our goal is simple: predict for each edge in a tree whether it is in a maximum parsimony tree or not. Our model consists of a recurrent neural network that learns features while traversing the input tree, which are used to classify the edge. The model makes high-quality predictions for this NP-complete problem on simulated and empirical datasets for trees of various sizes. We believe it is a stepping stone towards efficient phylogenetic inference using deep learning.

---

Phylogenetic inference is a formidable computational task. The number of phylogenetic trees that could explain the evolution of a set of sequences grows super-exponentially with the number of sequences. Additionally, the space of all phylogenetic trees is a complex structure (Holmes, 2003) with a complex optimality landscape (Maddison, 1991; Sanderson et al., 2011; Whidden and Matsen, 2015; Barker et al., 2025; Gao et al., 2025). Despite this, current algorithms are able to infer phylogenetic trees for thousands or even millions of sequences.

These algorithms are enabled by two concepts: local search (via tree rearrangements) and dynamic programming (e.g. the Fitch (1971) and Felsenstein (1973) algorithms). Given a starting tree, e.g. computed using a distance-based method like Neighbor Joining (Saitou and Nei, 1987), tree search algorithms use tree rearrangements to obtain all neighboring trees of the current tree. For every neighboring tree they compute a score using dynamic programming, e.g. likelihood (Kozlov et al., 2019; Minh et al., 2020) or parsimony score (Fitch, 1971). By moving to the neighboring tree with the best score, the tree search eventually finds a tree that cannot be improved further by this procedure, a local optimum.

This procedure requires evaluating every move of a given type (typical in the maximum likelihood approach (Kozlov et al., 2019; Minh et al., 2020)) or randomly sampled moves (typical in the Bayesian approach). There are a linear number of such moves for small perturbations such as nearest-neighbor interchange and a quadratic number for subtree prune and regraft. Because most moves are catastrophic for the objective function, this raises the question: *is more intelligent move selection possible?* Existing methods involve either characterizing the set of good trees (Höhna and Drummond, 2012; Barker et al., 2025), evaluating a less expensive objective function (Stamatakis et al., 2005; Zhang et al., 2020), or using tree summary statistics (Azouri et al., 2021, 2024).

In this paper, we present a new deep learning approach to find regions in a tree that need improvement in a single pass. Unlike previous applications of deep learning, the algorithm ingests the output of a dynamic program on the sequence alignment, rather than the raw sequence alignment. The algorithm then learns features based on these processed versions of the sequence data, which it uses to classify whether an edge belongs in the tree or not. We call this approach DPVT– Deep neural networks for Phylogenetics Via Traversal. This model uses a recurrent neural network in the shape of an input tree to learn features for all edges that are then used to classify edges. Though similar types of graph shaped neural networks exist (Tai et al., 2015; Thost and Chen, 2020; Ren et al., 2021), we are not aware of a version like ours where a simple recurrent network is used with the goal of classifying edges. By design, our model does not require trees to be of fixed size in training and testing datasets, which is a common restriction in existing deep learning methods for phylogenetics.

Although we formulate DPVT as a general framework for classifying edges, in this paper we use it to predict which edges in a provided tree are suboptimal under the maximum parsimony criterion. We show that this problem is NP-complete, and thus captures a challenging aspect of phylogenetic inference.

We assess the performance of our model by answering the following questions:

1. Can our model predict whether edges are present in a maximum parsimony tree or not?
2. Is the model able to generalize from simulated training data to empirical testing data?
3. Does a different distribution of non-maximum parsimony edges in the trees in training and testing data influence model performance?

We train and test DPVT on both simulated data and empirical data. The model is able to make high-quality predictions on both simulated and empirical datasets, even if the data contains trees larger than those in the training set. Performance is influenced by the distribution of maximum parsimony edges, but this effect can be mitigated by the choice of training data. Our results suggest that this new type of deep learning model for phylogenetics is a promising basis for guiding tree search.

## Methods

We begin our Methods section with a general overview of the DPVT model. Following this, we give a detailed description of the problem we are aiming to solve, provide a technical description of our deep learning models (DPVT), and explain the data we use for training and testing our model.

### Overview of the DPVT Model

We introduce DPVT (Deep neural network for Phylogenetics Via Traversal), a novel deep learning model that predicts which edges in a candidate tree are present in a maximum parsimony tree for the provided alignment. Edges are identified by their split: the bipartition of the set of leaves that results from removing this edge from the tree. If the split induced by an edge is present in a maximum parsimony tree, we call this edge an *MP edge*, otherwise it is called a *non-MP edge*. Deciding whether an edge is an MP edge or non-MP edge is *NP*-hard (Theorem 2).

DPVT combines dynamic programming and deep learning to classify edges in a candidate tree as being MP edges or not. The input is a candidate tree with sequences on all nodes, and we assume that this tree is binary and rooted in a leaf. We use the Fitch algorithm (Fitch, 1971) to infer mutations along edges and encode these mutations as numerical vectors, so that the input to the actual DPVT model is a tree with numerical vectors on all edges, as depicted in Figure 1.

**Figure 1:**
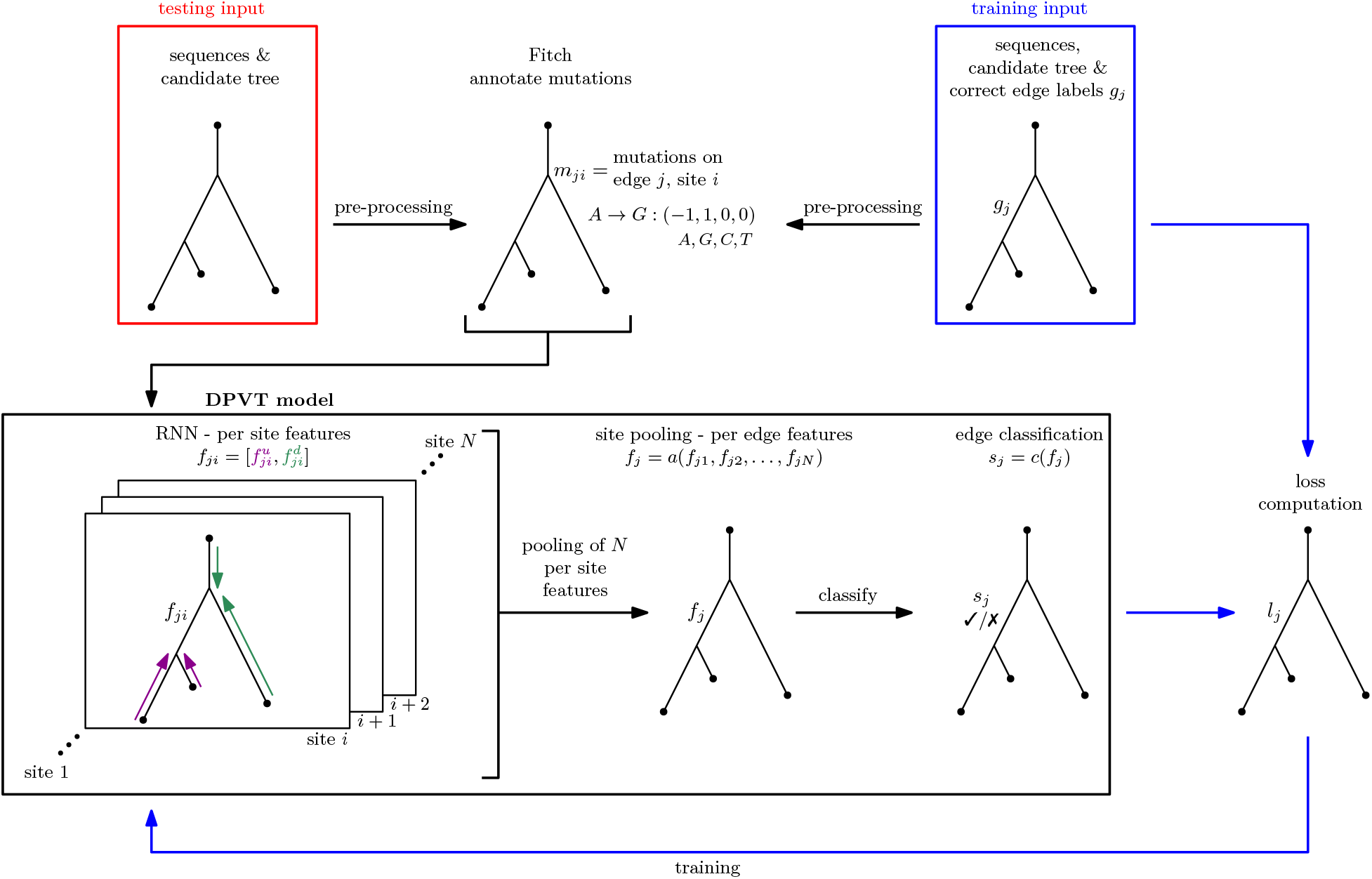
Overview of the DPVT model (in black box), pre-processing of input trees (top row) and training steps (blue arrows). When provided an input tree with sequences on all nodes, the pre-processing in the top row computes mutation encodings for each edge. The resulting tree with mutation encodings along edges is input for the two step DPVT model (RNN and site pooling step). The output of DPVT is a classification for each edge in the tree. For training, trees with correct edge labeling are used as input (top right blue box). After pre-processing, they are passed to the DPVT model, which computes edge predictions. Comparing those to the true labels yields a loss (bottom right tree) that is used to train the DPVT model.

To summarize, DPVT consists of two steps. The first step performs feature learning on each alignment site separately using a two-pass algorithm structurally similar to other such algorithms found in phylogenetics. However, rather than a fixed algorithm, this uses a trainable “recurrent unit” neural network to traverse the tree. The second step uses the per-site information on each edge to produce predictions with a second neural network. We now describe these steps in a little more detail.

During the first step, DPVT learns site-level features that are later used to decide whether an edge is present in a maximum parsimony tree. We use a recurrent neural network (RNN) that is adapted to the shape of the input tree. RNNs usually consist of a linear sequence of recurrent units and take sequential data like text or DNA sequences as input. They iterate through the input sequence and at each step feed the next element of the sequence into their recurrent unit, which contains a hidden state that holds information from previous steps. We do not provide a linear sequence as input, but a phylogenetic tree. Such tree-shaped neural networks have been used for other tasks (Tai et al., 2015; Ren et al., 2021).

In our RNN, each recurrent unit corresponds to an edge of the input tree and takes site-specific mutation encodings and previously-learned features of two adjacent edges as input. The order of the steps, and therefore recurrent units, is given by two tree traversals: In the first bottom-up (postorder) traversal, the input to the recurrent unit comes from adjacent edges on the non-root side of the current edge to learn its feature. In the second top-down (preorder) traversal, the information on the root side of an edge is used to compute its feature. This way our model uses all information on either side of an edge to learn the feature for this edge for a particular site.

The second step of DPVT is a pooling step where all per-site features for an edge are aggregated to one feature per edge. We compare three different models for this site pooling step: taking the average (average pooling) or the maximum (max pooling) of the learned features, or using the features as input to a transformer encoder (Vaswani et al., 2017) before averaging. After pooling the per-site features, we use a linear layer and a sigmoid activation function to classify each edge as an MP edge or not.

To train our model we require a dataset consisting of trees with correct edge labels indicating whether an edge is an MP edge or not. We provide a pipeline to generate such training data. Given a multiple sequence alignment, it runs larch (Barker et al., 2025) to compute a collection of maximum parsimony trees, and perturbs those trees to introduce non-MP edges.

To assess the quality of DPVT predictions, we compare its performance to that of a simple baseline model. This baseline model labels an edge as non-MP edge if there is a reversion on this edge, i.e. a mutation back to a character that has been present at the same site in an ancestral sequence.

Before explaining the DPVT model in detail, we introduce the MP Edge Problem, to which we apply the DPVT model.

### Complexity of the MP Edge Problem

The maximum parsimony principle for phylogenetic tree inference is based on the idea that mutations are rare. The optimal tree is assumed to be the one that minimizes the number of mutations along its edges (Fitch, 1971). Finding such a maximum parsimony tree is an *NP*-complete problem (Day and Sankoff, 1986). Here, we consider a slightly different but closely related problem: given a candidate tree, can we identify for each edge in this tree whether it is optimal according to the maximum parsimony principle?

We require some definitions to formally introduce this problem. Let *T* be a phylogenetic tree with leaf labels so that there is a bijection between these leaf labels and the sequences of a multiple sequence alignment *A*. For simplicity, we refer to the leaves of *T* using the corresponding sequences in *A*. The parsimony score ps*_A_*(*T*) of *T* is defined as the minimum number of mutations required for the sequences of *A* to fit on the tree *T*. A maximum parsimony tree is a tree that minimizes the parsimony score among all possible trees for *A*.

Instead of finding a maximum parsimony tree directly, we aim to predict for every edge in any given tree *T* whether it is present in a maximum parsimony tree for *A*. We identify an edge *e* by its split *S*_1_|*S*_2_, which is a bipartition of the set of leaves of a tree *T* into sets *S*_1_ and *S*_2_ so that removing *e* from *T* separates its leaf set into sets *S*_1_ and *S*_2_. There might be multiple maximum parsimony trees, and we want to predict whether an edge is in any of these trees or in none of them. We can phrase this problem as a decision problem:

**Problem 1** (MP Edge Problem). Let A be an alignment, S_1_|*S*_2_ *a split of A, and k an integer. Is there a tree T for A containing an edge inducing the split S*_1_|*S*_2_ *with* ps*_A_*(*T*) ≤ *k?*

**Theorem 1.** *The* MP Edge Problem *is NP-complete*.

The proof of Theorem 2 can be found in the supplement.

### DPVT for the MP Edge Problem

The DPVT model (Figure 1) takes a tree as input and uses its mutation history to make predictions for each edge. We use this model as a heuristic for the MP Edge Problem. In the following, we describe the individual components of the DPVT model and how we apply it to our problem, as well as how we generate training data.

### Pre-processing

We assume that input trees are binary and rooted in a leaf, and we know the mutation history: which mutations happen on each edge of the tree. We later explain how to generate trees of this format from a multiple sequence alignment. Because deep learning models require numerical vectors as input, we encode tree traversals and mutations along edges and sites as multidimensional vectors (“tensors”) and provide these two tensors as input to DPVT. We save the mutation encoding for an edge at its boundary node that is furthest from the root. This is made possible by the fact that each node in a tree (except for the root node) has exactly one incoming edge from the direction of the root.

For each tree, we store the information gathered in these traversals in tensors of dimension 2 ×(*n* − 3) × 3, where *n* is the number of leaves of the input tree. The first dimension is 2, which is the number of traversals that we perform during training: bottom-up and top-down traversals. The second dimension is *n* − 3, the number of internal edges of a binary tree rooted in a leaf. The third dimension is 3, as we save the indices [*i_x_, i_y_, i_v_*] of three nodes *x, y, v* for each edge we visit. For the bottom-up traversal, *v* is the node bounding the current edge on its non-root side, and *x* and *y* are its children. In the top-down traversal, *v* is the node bounding the current edge on its non-root side, *x* is its parent, and *y* its sibling. We index nodes according to a preorder traversal and generate the traversal data structure for each tree by performing a postorder traversal followed by a preorder traversal. We skip edges that are incident to leaves (including the root leaf) as these edges are present in every tree on the same leaf set and hence in every maximum parsimony tree.

While performing the preorder traversal, we also compute the mutation encodings by iterating over all sites for each edge, thereby creating a tensor of dimension (2*n* − 3) × *N* × 4, where *N* is the number of sites. Unlike the traversal tensor, this mutation tensor includes all 2*n* − 3 non-root edges (internal and pendant), since the recurrent unit at an internal edge adjacent to leaves still requires the mutation encodings of those pendant children. This tensor contains for each edge (*k*, first dimension) and each site (*i*, second dimension) an entry *m_ki_* = [*µ_A_, µ_G_, µ_C_, µ_T_*] describing the mutation at this edge and site. If there is a mutation from nucleotide *x* to *y* (in the direction away from the root) with *x ≠ y*, we set *µ_x_* = −1, *µ_y_* = 1, and *µ_z_* = 0 for *z ≠ x, y*. If the edge has no mutation at this site, the entry is the zero vector [**0**] = [0, 0, 0, 0]. An example is displayed in Figure 2.

**Figure 2:**
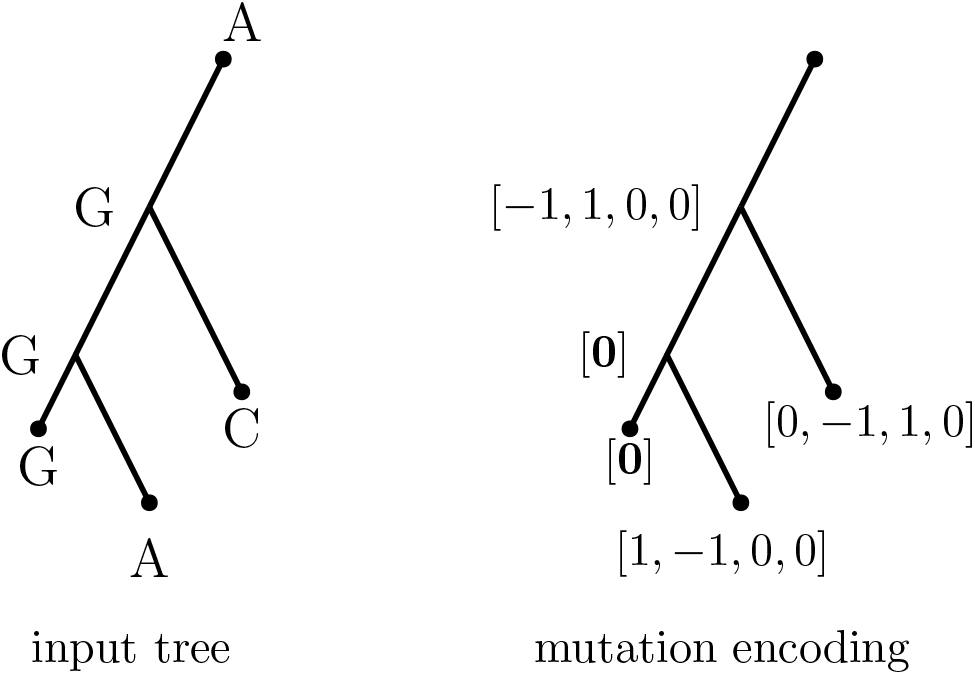
Input tree with sequences at one site (left) and mutation encodings (right). [**0**] represents the zero vector [0, 0, 0, 0], i.e. no mutations.

### The DPVT Model

#### Step 1: Tree Traversal

In the first step of our model we apply a neural network to learn features from the tree structure and mutations along its edges. We apply the same neural network independently to all sites of the sequence alignment, so that the learned features encode information for each site and edge. This neural network can be thought of as a recurrent neural network (RNN) that follows the shape of the tree, where the recurrent unit is used to compute a feature for each edge and site of the tree. We therefore traverse the input tree and at each edge we visit, the already learned features and mutation encodings of the two edges leading to this edge are the input to our recurrent unit to compute a feature for the edge (see Figure 3). By doing this in two traversals, once from the leaves to the root (“upward”, blue arrows in Figure 3) and then from the root to the leaves (“downward”, red arrows in Figure 3), we use all information on either side of an edge to learn these features. We concatenate the features learned in the two traversals to one *per-site-and-edge feature*, as we consider each site separately in this step.

**Figure 3:**
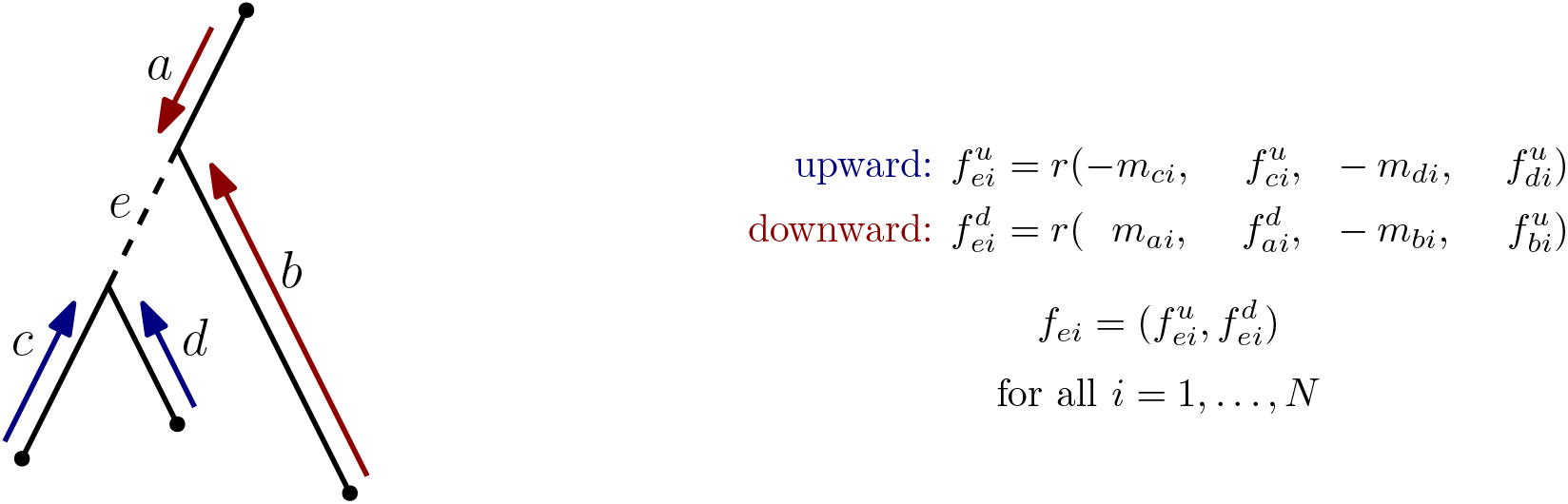
Illustration of the RNN used in the first step of DPVT. Tree with edge labels shown on the left and computation of per-site-and-edge feature *f_ei_* for edge *e* and site *i* shown on the right, using the recurrent unit *r*. Mutations at edge *k* and site *i* are represented by mutation vectors *m_ki_*.

The recurrent unit of the RNN we use in this step of DPVT is a two-layer feed-forward network with ReLU activation function between the layers. The hidden layer dimension is set as a hyperparameter. For each internal edge we visit during the traversal, we iterate over all sites of the sequences, yielding *N* per-site features for that edge. We initialize features of pendant edges, i.e. edges incident to leaves, as zero vectors. The features learned during upward and downward traversal are kept separate while traversing, but are concatenated afterwards. For example, for an edge *e* and site *i* we learn feature 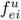 in the upward traversal and 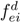 in the downward traversal and concatenate them afterwards to one feature 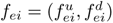.

In the first traversal, a postorder traversal upward from leaves to root, the input to the recurrent unit *r* are mutation encodings *m_ci_, m_di_* and already learned features *f_ci_, f_di_* of the two edges *c* and *d* that are adjacent to the current edge *e* and pointing away from the root (see Figure 3). We multiply the mutation features of the two input edges by −1, as they are encoded in the direction away from the root, whereas we are interested in using all information in the direction toward the root to learn features in this upward traversal.

In the second traversal, a preorder traversal from root to leaves, mutation encodings *m_ai_, m_bi_* and features 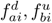 corresponding to the two edges *a* and *b* on the root side of the current edge *e* are used as input to the recurrent unit *r*. Assuming that *a* is the edge on the path from the current edge to the root, we are using the upwards traversal feature for edge *b*, as we want to use all information collected in the subtree below *b* when computing the feature for *e*. We also multiply *b*’s mutation encoding by −1, as this edge is pointing upward toward *e*, while the mutation encoding assumes the edge is pointing away from the root. Due to these directionality considerations, the choice of leaf for the root of the tree influences how features are computed, which makes DPVT effectively a method for rooted trees. Note that the feature for an edge *e* is computed only from information on either side of *e*, not from *e*’s own mutation encoding. The detailed formula for computing features is given on the left of Figure 3.

#### Step 2: Site Pooling

After the traversal step, every edge *e* has *N* per-site features *f_e_*_1_*, f_e_*_2_*, …, f_eN_*. Each feature is the concatenation of the upward and downward features for that edge and site, as described in the previous step: 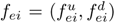. In the second step of our model we use pooling to aggregate these features to one feature per edge. We use one of three pooling functions *a* to compute the feature *f_e_* = *a*(*f_e_*_1_*, f_e_*_2_*, …, f_eN_*) for edge *e* (see Figure 1):

- averaging
- maximizing
- transformer encoder followed by averaging

The simplest way to aggregate all per-site-and-edge features to one per-edge feature is to use average pooling or max pooling, taking the average or maximum of the per-site-and-edge features. For the latter, we maximize over all sites by taking for each position *k* of the feature vectors the maximum entry over all per-site features for this edge.

Our third model uses a transformer encoder before average pooling to aggregate features. The input to the transformer is a tensor of *E* × *N* × *d*, where *E* is the number of internal edges in our tree, *N* is the number of sites, and *d* is the feature dimension. The output is a tensor of the same size, which we then average over all *N* sites to receive an *E* × *d* tensor containing one feature per edge. After aggregating to one per-edge feature, we use a linear layer and a sigmoid activation function to classify each edge as being an MP edge or not.

### Baseline Model

To assess the quality of the predictions of DPVT for the MP Edge Problem, we compare it to a simple baseline model. This baseline model takes the same input as DPVT: a candidate tree with sequences on all nodes. An edge is predicted to be a non-MP edge if at any site there is a reversion on this edge, i.e. a mutation back to a state present at this site in an ancestral sequence. All other edges are labeled MP edges.

### Training

For training, we use the Adam optimizer (Kingma and Ba, 2017) and binary cross entropy as the loss function. As the transformer encoder expects a fixed input size, we pad sequence lengths to get the same number of sites for all sequences. This padding is masked in the transformer encoder and when computing loss. We also mask edges incident to leaves (including the root) when computing loss, as the splits induced by these edges are present in all trees, and therefore will always be MP edges. During training, we use a validation set to check whether our model is overfitting and use early stopping when validation loss does not decrease within five epochs. Our hyperparameters are learning rate, batch size, number of epochs, feature length, and dimension of the hidden layer of the RNN. After some initial hyperparameter search with Optuna (Akiba et al., 2019) on simulated data with 15 leaves, we fix these hyperparameters to: learning rate 5 × 10*^−^*^5^, batch size 4, number of epochs 200, feature length 64, and dimension of hidden layer 256.

### Training and Testing Data Generation

To train our classifier, we generate phylogenetic trees with edges labeled as MP edges or non-MP edges. In the following we explain how we compute maximum parsimony trees for simulated or empirical alignments with larch (Barker et al., 2025), and how we follow this up with introducing non-MP edges by perturbing these maximum parsimony trees.

#### Generating MP Trees

We provide a pipeline to create training data from (empirical or simulated) alignments and produce trees with edge labels, indicating whether each edge is an MP edge or not for the provided alignment. Before inferring maximum parsimony trees, we pre-process alignments by deleting all sites that are non-informative for maximum parsimony, i.e. sites that have the same nucleotide for all sequences and those where only one nucleotide differs from all others. Additionally, we remove sites containing gaps or ambiguous characters, and duplicate sequences.

When simulating alignments, we initially simulate alignments twice as long as the target length and with five more sequences than specified. This is to make sure that the alignment dimensions are not below the target size after removing non-informative sites and duplicate sequences. We remove excess sites and sequences after pre-processing to cut down to the target size and re-run the process if the target size cannot be achieved.

Once alignments are pre-processed, we use the software package larch (Barker et al., 2025) to generate a history DAG, a data structure containing a collection of maximum parsimony trees for each alignment. The goal of larch is to find the entire ensemble of parsimony optimal histories, and it has shown to perform well at this task (Barker et al., 2025). For our work we will assume that the resulting history DAG contains all maximum parsimony topologies. From this history DAG we randomly choose a maximum of 200 unique tree topologies per alignment, which we then perturb (as described below) to introduce non-MP edges.

#### Tree Perturbations

To generate non-MP edges in the maximum parsimony trees returned by larch, we use two different types of tree perturbations. We either perform a subtree prune and regraft (SPR) move or a *random bounded-depth subtree replacement*. A bounded-depth subtree of a tree is a subgraph that is given by a node *v* and depth *d* so that the subgraph contains *v* and all nodes reachable from *v* by traversing at most *d* edges in direction away from the root. For a random bounded-depth subtree replacement (see Figure 4), we assume that a depth *d* is given and pick a random node *v* with height greater than or equal to *d*, i.e. there is at least one leaf that is connected to *v* by at least *d* edges. We then remove the bounded-depth subtree with depth *d* that is rooted in *v*, while keeping all nodes that are leaves of this bounded-depth subtree. This gives us a disconnected graph *G* where *v* is a leaf and there are *x* nodes whose incoming edges have been removed. We generate a random tree with *x* leaves from a uniform distribution, make *v* its root, and replace its leaves by the *x* nodes without incoming edges from the graph *G* in random order. We choose the depth *d* to be half of the total tree depth, which is the maximum number of edges between the root of the tree to any of its leaves.

**Figure 4:**
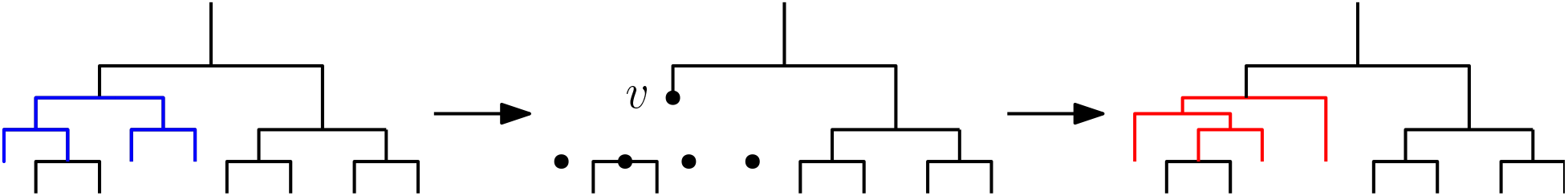
Introducing non-MP edges by a random bounded-depth subtree replacement. This replaces the subgraph in blue on the left by the one in red on the right via the intermediate graph *G* in the middle.

For either type of perturbation, we can use the history DAG returned by larch to determine whether the newly introduced edges are present in a maximum parsimony tree or not. This allows us to generate 0*/*1 labels for all edges, which indicate whether an edge is present in a maximum parsimony tree (MP edge, 0) or not (non-MP edge, 1). Note that by “present in a maximum parsimony tree” we mean that the split induced by the edge is also induced by an edge in a maximum parsimony tree. For random bounded-depth subtree replacements we repeat the perturbation procedure until at least a third of all edges are non-MP edges.

Input trees to the DPVT model are expected to have sequences on all nodes of the tree, including internal nodes, from which we are able to label edges with mutations. We therefore run the Fitch algorithm to infer internal node sequences for the trees after perturbation. Whenever there are multiple options for a character at a site in the downward phase of the Fitch algorithm, a random one is chosen. The mutation history is a latent variable that we cannot observe directly, and these random tie-breaking choices effectively sample from the set of equally parsimonious histories. Training across many such reconstructions thus exposes the model to a distribution over plausible histories rather than a single deterministic one, which we expect to encourage robustness to this ambiguity.

### Data

We use simulated data to train our models and simulated and empirical alignments to test them. The alignments are used as input to the data generation pipeline described above, which returns trees with correct labels of non-MP and MP edges to train and test our models.

We simulate Yule-Harding trees and alignments for these under a Jukes-Cantor model using the alisim package in IQ-TREE2 (Ly-Trong et al., 2023). For training, we use two sets of 500 alignments: one on 25 and one on 50 sequences, both with 100 sites. Testing data is generated independently of training data using the same pipeline, resulting in 200 alignments with 25 or 50 sequences and 100 sites. To test our models on empirical datasets, we use data from three different sources:

- endemic human virus data (Kistler and Bedford, 2023): influenza C segments M, NS, and PB2, and rotavirus A segment 11,
- orthologous mammalian markers from the OrthoMaM database (Allio et al., 2024), and
- coding DNA sequence of a variety of families of homologous protein domains from the PANDIT database (Whelan et al., 2006).

We use a filtering step for the alignments from the OrthoMaM and PANDIT databases to generate training and testing sets of good quality and feasible size for our DPVT models. We first remove all sequences where more than 20% of characters are gaps or ambiguous characters. Then we remove all sites containing gaps or ambiguous characters. Sites that are uninformative to maximum parsimony inference, i.e. those that contain the same character for all sequences or those where only one sequence contains a character different from that in the other sequences, are removed after this. For the PANDIT dataset we only keep alignments where at least 80% of sites of the original alignment are present after this processing. The resulting alignments are also split into 80% training and 20% testing sets. For OrthoMaM we set the site filtering threshold to 50%, as we otherwise discard too many datasets. We split the resulting alignments into training and testing sets and then randomly draw 1, 000 trees from the training set and 200 from the testing set. In contrast to this, we generate the testing data for each of the endemic human viruses from just one alignment. Testing on the data from multiple alignments from OrthoMaM and PANDIT databases allows us to explore how well the model generalizes, while testing on specific alignments for viral datasets gives us an idea of how well the model works for each of these particular alignments.

### Implementation

An implementation of our models is available on GitHub: https://github.com/matsengrp/dpvt. We additionally provide the training data generation workflow as well as an implementation of a pipeline for training those models: https://github.com/matsengrp/dpvt-experiments-1. Our implementation is written in Python with coding support from Claude (Anthropic, 2025) and uses PyTorch-Lightning and torchmetrics (Falcon and The PyTorch Lightning team, 2019) for implementing the neural network models, Optuna (Akiba et al., 2019) for hyperparameter optimization, ete3 (Huerta-Cepas et al., 2016), historydag (Dumm et al., 2023), and alisim (Ly-Trong et al., 2023) for generating and manipulating trees, and Snakemake (Mölder et al., 2021) for implementing the workflows for data generation, training, and testing. Other auxiliary packages used are: BioPython (Cock et al., 2009), black (Langa and contributors to Black, 2025), matplotlib (Hunter, 2007), numpy (Harris et al., 2020), pandas (McKinney, 2010; pandas development team, 2025), pytest (Krekel et al., 2004), scikit-learn (Pedregosa et al., 2011), tensorboard (Abadi et al., 2016), IQ-TREE (Minh et al., 2020), usher (Turakhia et al., 2021), tbparse (Sun, 2025), and seaborn (Waskom, 2021).

## Results

### DPVT Models Make Accurate Predictions for Simulated Data

To evaluate the quality of our model’s predictions, we first train and test on simulated data. We simulate sequence data under a Jukes-Cantor model using the alisim package in IQ-TREE2 (Ly-Trong et al., 2023). To generate training data, we simulate two sets of 500 alignments: one on 25 and one on 50 sequences, both with 100 sites. We separately simulate two testing sets of 200 alignments with the same properties. We compute a collection of maximum parsimony trees for these alignments using larch (Barker et al., 2025) and take a maximum of 200 of these trees. To introduce non-MP edges, we apply 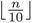 Subtree Prune and Regraft (SPR) moves to each tree, where *n* is the number of leaves of a tree. The fraction of non-MP edges in the resulting trees is displayed in Figure S7. To evaluate the performance of our model, we compute the area under the receiver operating characteristic curve (AUROC) for our model predictions (Figure 5). We display three different types of model along the y-axis that differ by the method used in the second step of DPVT, where the per-site features are pooled to one feature per edge (averaging, maximizing, and transformer encoder).

**Figure 5:**
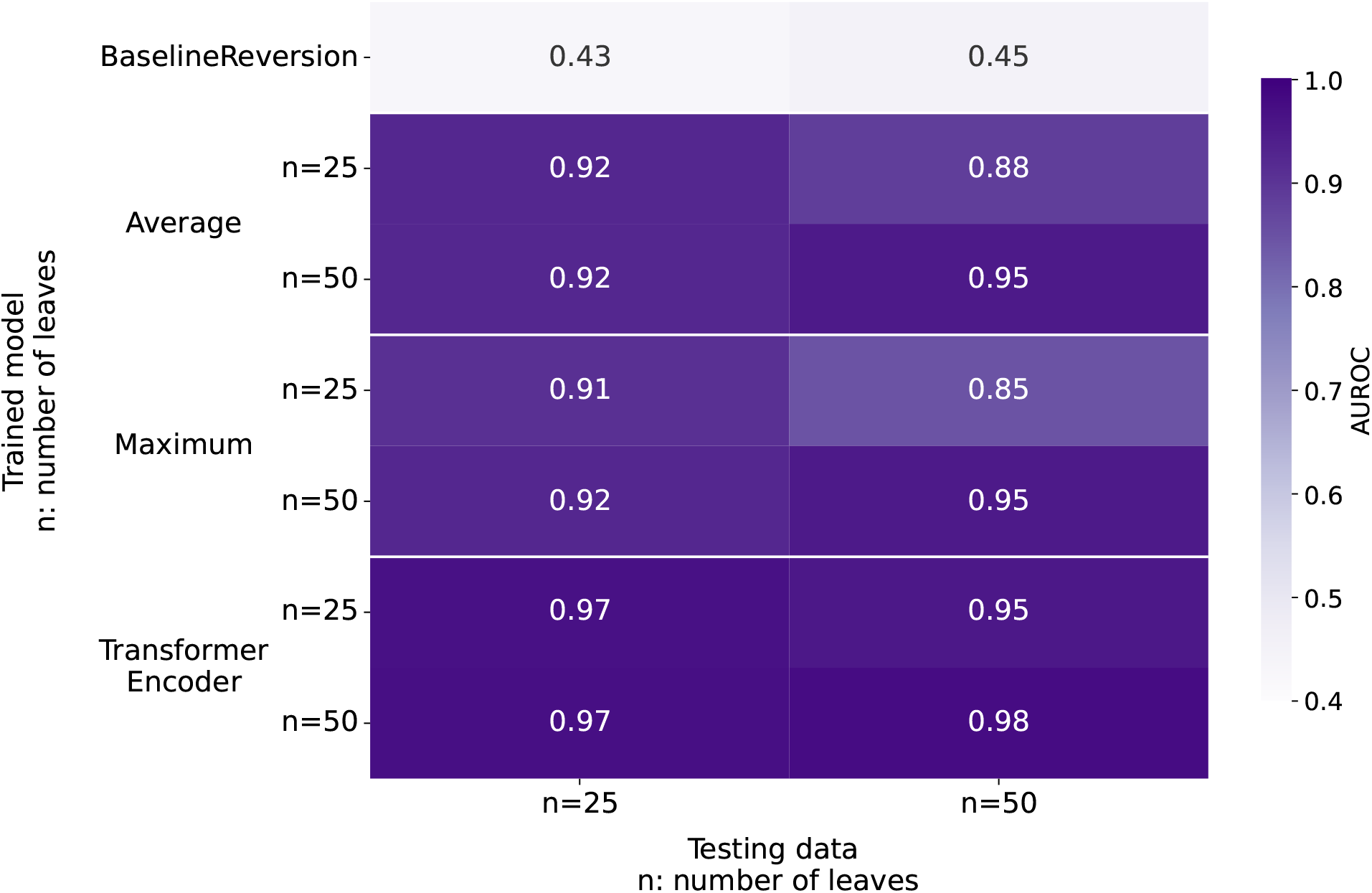
Model performance on simulated data for models trained on simulated data. AUROC of testing data on 25 and 50 leaves (x-axis) for different DPVT models trained on data on 25 and 50 leaves (y-axis). Three different site pooling methods for DPVT are displayed on the y-axis: averaging, maximizing, or applying a transformer encoder before averaging. The top row corresponds to the performance of the baseline model, which does not need to be trained.

All DPVT models show high AUROC values, especially in comparison to the baseline model that has an AUROC of 0.43 and 0.45 for the 25 and 50 sequence test set, respectively. The transformer encoder pooling model performs best with AUROC values between 0.95 and 0.98, while all other models perform only marginally worse. All models show their best performance when trained and tested on trees with the same number of leaves, and achieve highest AUROC when trained and tested on 50 leaf datasets.

Models trained on simulated data perform slightly worse when tested on trees with a different size than training trees. For example, for the average pooling model trained on 25 leaves we observe an AUROC of 0.92 when testing on 25 leaf trees and an AUROC of 0.88 when testing on 50 leaf trees. This difference is only marginal, though, indicating that the models are able to generalize from smaller to larger trees.

### Models Trained on Simulated Data can Generalize to Empirical Data

To determine how well DPVT generalizes from simulated to empirical data, we test the models trained on simulated datasets as described above on empirical data (Figure 6). We generate testing data for empirical alignments of viral datasets (influenza C segments M, NS, and PB2 and rotavirus A segment 11 from Kistler and Bedford (2023)) and datasets taken from two different databases: OrthoMaM (Allio et al., 2024) and PANDIT (Whelan et al., 2006). For each of the viral datasets, we have one alignment to generate testing data, while we take multiple alignments from the databases to generate training and testing datasets. More details on the datasets can be found in Methods. Due to memory constraints when using the transformer encoder pooling in DPVT, we only use average and maximum pooling for these datasets, which have hundreds of sites.

**Figure 6:**
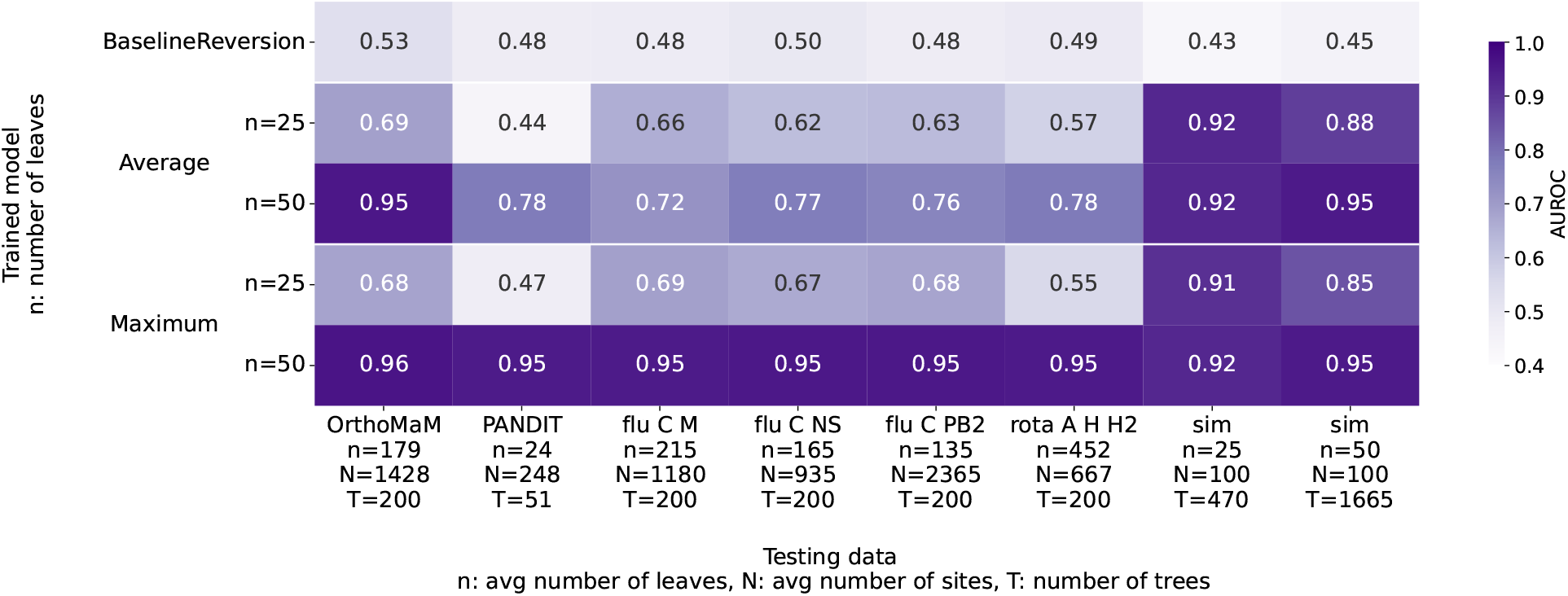
Model performance on empirical and simulated data for models trained on simulated data. The y-axis displays different models: the baseline model and average and maximum pooling DPVT models, which are trained on simulated datasets with varying number of sequences, as shown on the y-axis labels. The x-axis displays the different testing datasets, including the number of sequences and sites of the corresponding sequence alignment.

When testing our models trained on simulated data on empirical data, we find that the maximum pooling model trained on the 50 leaf dataset performs exceptionally well with AUROC values of 0.92 and above. The same is true for the average pooling model on 50 leaves when tested on the OrthoMaM testing set. All other models show worse performance on empirical data than on simulated data with AUROCs on empirical data between 0.44 and 0.69 when training on *n* = 25 leaf data and between 0.72 and 0.78 on *n* = 50 leaf data. This suggests overfitting of these models on simulated data, especially when trained on 25 leaf trees.

We also trained our DPVT models on an empirical dataset (Figure 7). To create training data, we filtered and subsampled data from the OrthoMaM database, which we describe in more detail in Methods. Most importantly, we made sure that different alignments are used for generating training data and testing data. We observe that models trained on empirical data perform very well on all empirical testing datasets and are comparable to the maximum pooling model trained on simulated data with *n* = 50 leaves. The worst performance among all empirical datasets shows on the PANDIT data, where the AUROC is 0.78 for average pooling and 0.81 for maximum pooling. The reason for this might be the smaller size of the trees in this dataset compared to that in the training set. The average number of leaves *n* in the training data is 152, while the PANDIT trees average *n* = 24 leaves. For all other empirical datasets the AUROC is at or above 0.92. When testing on simulated data, however, we observe lower AUROCs, which again might be caused by our simulated trees having fewer leaves (*n* = 25, 50) than the training data. Testing on datasets with larger trees and more sites than the training data does not seem to be a problem to our model, as it still performs well for the rotavirus dataset that has 452 sequences and 667 sites.

**Figure 7:**
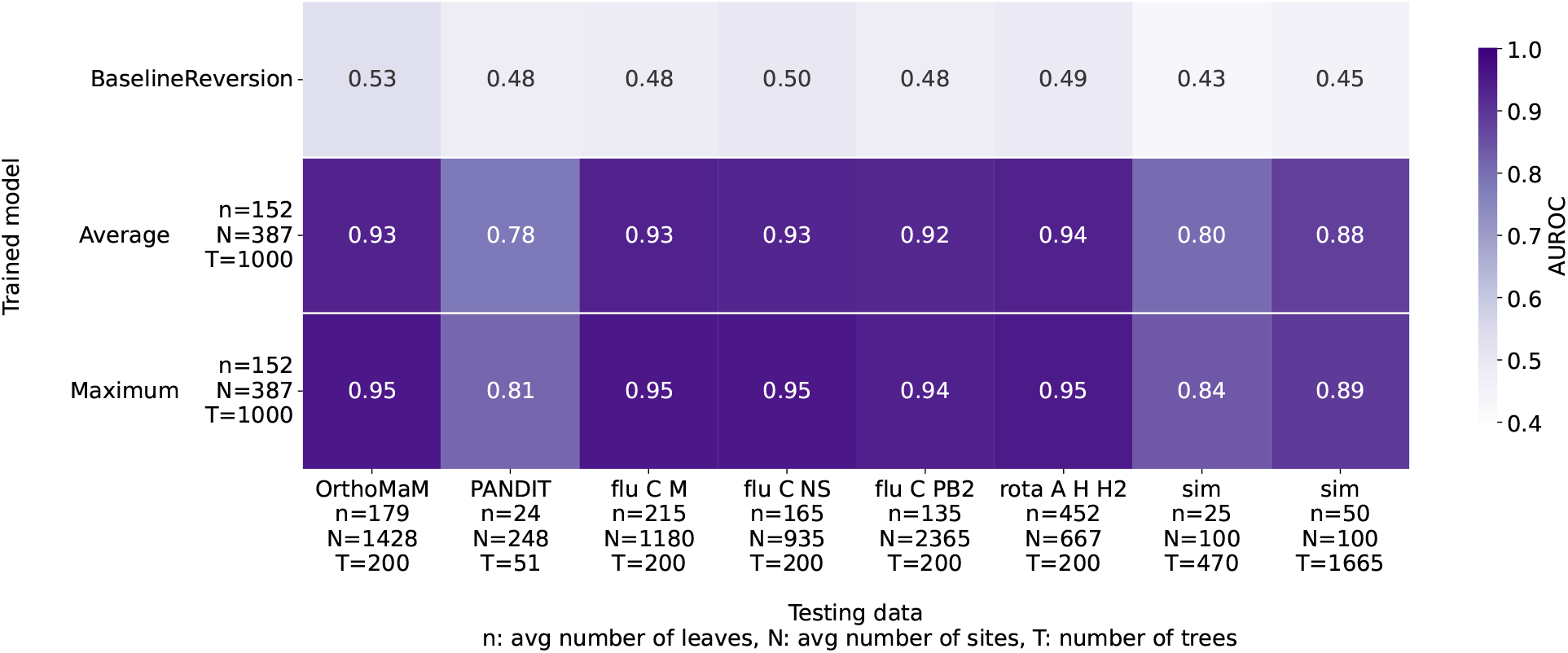
Model performance on simulated and empirical data for models trained on empirical data from the OrthoMaM database. The y-axis displays different models: the baseline model and average and maximum pooling DPVT models. The x-axis displays the different testing datasets, including the number of sequences and sites of the corresponding sequence alignment.

As our testing sets are imbalanced in the number of non-MP edges (see Figure S7), we also computed precision-recall curves to compare model performances. They confirm the similar performance of the maximum pooling model trained on simulated data on 50 leaves and the models trained on empirical data (Figures S1, S2, S3, S4).

### Prediction Quality is Sensitive to the Distribution of Non-MP Edges

Depending on the source of candidate trees used as input to DPVT, the distribution of non-MP edges might differ. To see whether DPVT model performance is influenced by different distributions of non-MP edges in training and testing trees, we generate different datasets by introducing non-MP edges using one of two types of tree perturbations on our simulated alignments: SPR moves (as in the previous sections) or random bounded-depth subtree replacements. For the latter, we take a random node in an MP tree and remove all paths of random length *s* that start at this node and point away from the root. We then introduce a new graph structure replacing the depth *s* subtree that was removed with a random rooted binary tree. This procedure is repeated until at least a third of all edges are non-MP edges. We describe this perturbation in more detail in our Methods. Different perturbation methods lead to different distributions of non-MP edges, visible in the number of non-MP edges introduced (Figure S7) and the length of the longest path of non-MP edges within a tree (Figure S8).

We train our models on three training sets generated from simulated data with 50 sequences: one where SPR moves introduce non-MP edges, one with random bounded-depth subtree replacements, and a third that is the union of these two sets. We test these models on datasets generated from different simulated alignments, using the same three perturbation approaches (Figure 8). Different perturbation methods in training and testing data generation result in worse model performance, having AUROC values between 0.63 and 0.79, compared to values of 0.93 and above when the same perturbation method is used. When training on the union of SPR and random bounded-depth subtree replacement datasets, we see better performance, with all AUROC values at or above 0.92 even when non-MP edges are introduced in different ways in training and testing data. The transformer encoder model shows especially good performance, with AUROC values of 0.98 when trained on the mixed perturbation dataset.

**Figure 8:**
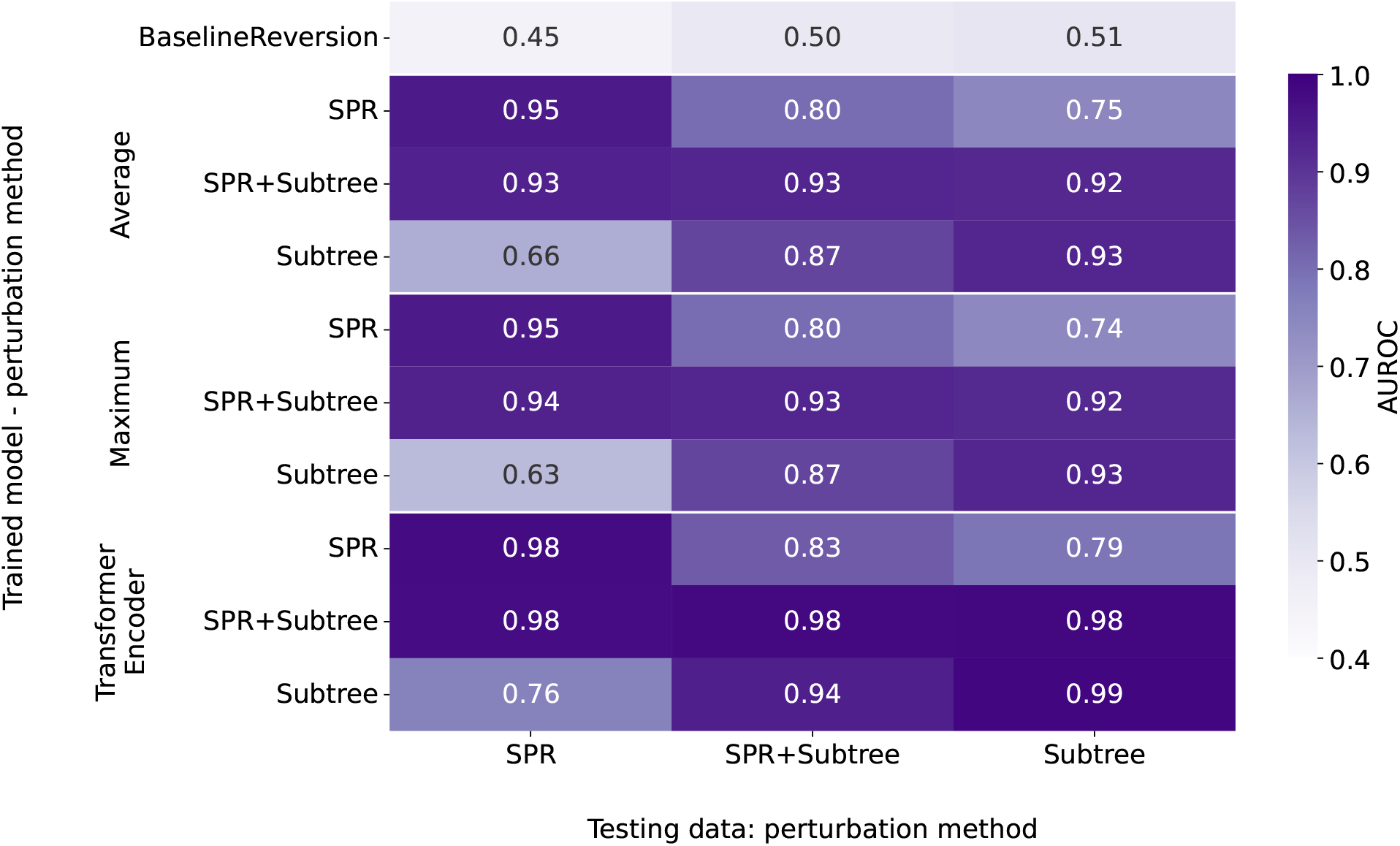
Training and testing on simulated data with *n* = 50 leaves and *N* = 100 sites. We use three types of tree perturbation to introduce non-MP edges in our training sets (y-axis): SPR moves (“SPR”), random bounded-depth subtree replacements (“Subtree”), or both (“SPR+Subtree”). Similarly, we use three testing datasets with these three tree perturbation methods. Top row shows performance of baseline model that labels edges with reversions as non-MP edges.

### Training and Testing Times

One great benefit of deep learning models is that once they are trained, they can make predictions very quickly. Our models are trained and tested on a single NVIDIA A100 GPU (80GB PCIe). All training times are shown in Figure S5. Training on simulated data with 25 leaf trees takes one and a half to three hours, with the transformer encoder model taking the shortest time. For the simulated data on 50 leaves, the transformer encoder and the average pooling model train for five and a half hours while the maximum pooling needs ten hours. The shorter training times of some models can be explained by training being stopped early, which happens when the validation loss no longer declines, indicating overfitting to the training data.

For the empirical data from the OrthoMaM database the training times are much longer than those for our simulated datasets, as they contain trees with more leaves and longer sequences. The average pooling model requires 44 hours and the maximum pooling model 52.5 hours.

Time required for testing our models depends on the size of the trees in the testing data. In Figure S6 we display the average wall time required for testing our model per tree for each dataset. We see that more sequences and sites in a dataset lead to longer times when testing the DPVT models. We observe the fastest testing times for simulated trees with about 0.025 seconds per tree while the rotavirus trees require the longest time for testing with about 0.35 seconds per tree.

## Discussion

In this paper we present DPVT, a novel approach for using deep learning in phylogenetics. This model learns features for all edges in a provided tree to classify edges. Here, we use DPVT to predict whether edges in a given tree are present in a maximum parsimony tree or not. Our model uses the reconstructed mutation history, i.e. mutations inferred along edges in the tree, to learn edge features. The main component of this model is a recurrent neural network that follows the shape of the input tree. This network architecture is a type of graph neural network (Zhou et al., 2020; Thost and Chen, 2020; Corso et al., 2024) in the shape of a tree, which gives us a partial ordering of nodes that is followed during training. This is similar to the directed acyclic graph neural networks proposed by Thost and Chen (2020), or tree shaped networks (Tai et al., 2015; Ren et al., 2021), where message passing follows the partial order provided by a directed graph. Unlike most applications of these types of networks, we are only interested in features for edges in our tree, and use fixed mutation encodings along edges combined with previously computed features as input to our recurrent units.

One benefit of DPVT compared to most existing deep learning models in phylogenetics (Mo et al., 2024) is that the design of the RNN in the traversal step allows using input trees with varying numbers of leaves. Because the model considers all sites separately in this step, a pooling step is used to summarize all per-site features to one feature per edge before classification. We provide three different options for pooling: maximum and average pooling and applying a transformer encoder before taking the average. With this design, DPVT is a flexible model that can be adapted to different applications, although we focus on the MP Edge Problem in this paper.

Our results show that DPVT makes high-quality predictions. When training on simulated data, we observe that the model using a transformer encoder outperforms simpler pooling mechanisms (averaging and maximizing), but differences in performance are minor. Restricting the size of training trees to *n* = 25 leaves for simulated data shows much worse performance than training on larger trees with *n* = 50 leaves. While this difference could be at least partially attributed to the choice of models in simulations, we focused on using larger trees and empirical data for training as it clearly shows superior performance. For the latter, the maximum pooling DPVT model performs exceptionally well with AUROCs of 0.92 and higher for all testing data, indicating that it is able to generalize to empirical datasets. The performance of DPVT models trained on empirical data is comparable to that of the maximum pooling model trained on simulated data with *n* = 50 leaves. This similarity in performance between the model trained on simulated data and the ones trained on empirical data might be caused by the simplicity of the maximum parsimony criterion, which is only interested in the number of mutations required by the tree.

We also find that models that are trained on sufficiently large trees (*n* = 50 leaves for simulated data, or the larger empirical trees) generalize to larger datasets, both in the number of sequences and the length of sequences. Different distributions of non-MP edges in training and testing data do affect performance, which can be mitigated by the choice of training data. This suggests that a good choice of training data or fine-tuning of the model is important to achieve good model performance. DPVT models trained on empirical data and tested on empirical data from different sources perform very well.

The DPVT approach presented here is distinct from existing neural-network approaches for phylogenetic inference. The first approaches leveraging deep learning for phylogenetic inference have treated input sequences like images and phylogenetic inference as a classification problem (Suvorov et al., 2020; Zou et al., 2020; Wang et al., 2023). To limit the number of classes, these methods infer quartets, i.e. phylogenies for subsets of four sequences (Suvorov et al., 2020; Zou et al., 2020). The inferred quartets can then be merged into one tree containing all given sequences (Wang et al., 2023). However, this approach is only more time efficient than standard likelihood methods for trees with fewer than 40 sequences. To overcome this issue, machine learning methods have been developed to guide tree moves in the maximum likelihood framework (Azouri et al., 2021, 2024; Ly-Trong et al., 2024), allowing a single improving move to be selected without comparing all neighboring trees. Currently, however, none of these methods have both high accuracy and efficiency for trees with hundreds of leaves.

An alternative deep learning approach introduced in the literature is the inference of distance matrices from multiple sequence alignments, which can then be used as input to distance-based methods to infer phylogenies (Nesterenko et al., 2025). The accuracy of this approach depends on how similar training and testing data are and decreases with increasing number of sequences. Memory requirements are very high for larger trees, reducing the applicability of this approach for large datasets, such as one encounters when studying viruses. To directly infer a tree from sequences, phyloGAN (Smith and Hahn, 2023) uses generative adversarial networks to construct a phylogeny for a dataset, with the disadvantage of needing to be newly trained for every input alignment.

Though our model is very good at making accurate predictions about whether edges are MP edges or not, there is room for improvement, especially when it comes to memory and time efficiency. We were not able to train the transformer encoder model on some of our data due to memory restrictions. Another limitation of this model so far is its restriction to maximum parsimony inference.

In the future, we will work on a more memory efficient version of our models, especially the transformer encoder model, to train and test on larger datasets. This could for example be done by parallelizing parts of the tree traversal step and using more efficient data structures. We are also curious to see to what extent each choice of model design influences performance. For example, is randomly rooting our input trees more effective than systematically rooting them? And what is the minimum size of a neural network to effectively predict non-MP edges? Is it possible to explain the reasoning of the model’s predictions?

The DPVT model itself is very general: it uses information collected during a tree traversal to make predictions for edges in the tree. It can therefore be applied to any task that requires per-edge predictions. The application motivating our work here is tree search. If we can assess the quality of edges using DPVT, we can use these predictions to guide tree moves, especially for maximum parsimony inference. Recent work on maximum parsimony inference has shown that in the densely-sampled regime, especially for viral data where many samples with few mutations are available, many maximum parsimony trees can be compactly captured as a collection in a data structure referred to as a “history DAG” (Barker et al., 2025). We think that DPVT could be extended to work on this data structure to help explore the maximum parsimony landscape more efficiently. An extension of DPVT to maximum likelihood inference also seems possible, but requires larger modifications of the model to incorporate partial likelihoods. As Bayesian inference methods are those that struggle most with large, and especially densely sampled, datasets (Gao et al., 2025), maximum parsimony principles have been used to design more efficient proposals in the Bayesian inference framework (Zhang et al., 2020; Bouckaert et al., 2025). If instead of classifying edges, we compute weights for edges to indicate how likely they are to be MP edges, we could use such weights to define proposal distributions for Bayesian inference via MCMC, similar to how it is done by Zhang et al. (2020) and Bouckaert et al. (2025).

## Data Availability

All datasets used in this paper can be accessed through DRYAD: https://doi.org/10.5061/dryad.0zpc867bp.

## Funding

This work was supported by NIH grant R01-AI162611. Scientific Computing Infrastructure at Fred Hutch is funded by ORIP grant S10OD028685. Frederick Matsen is an investigator of the Howard Hughes Medical Institute.

## Acknowledgments

The authors thank Marc Suchard for feedback on earlier versions of this project.

## Supplementary Materials

**Figure S1:**
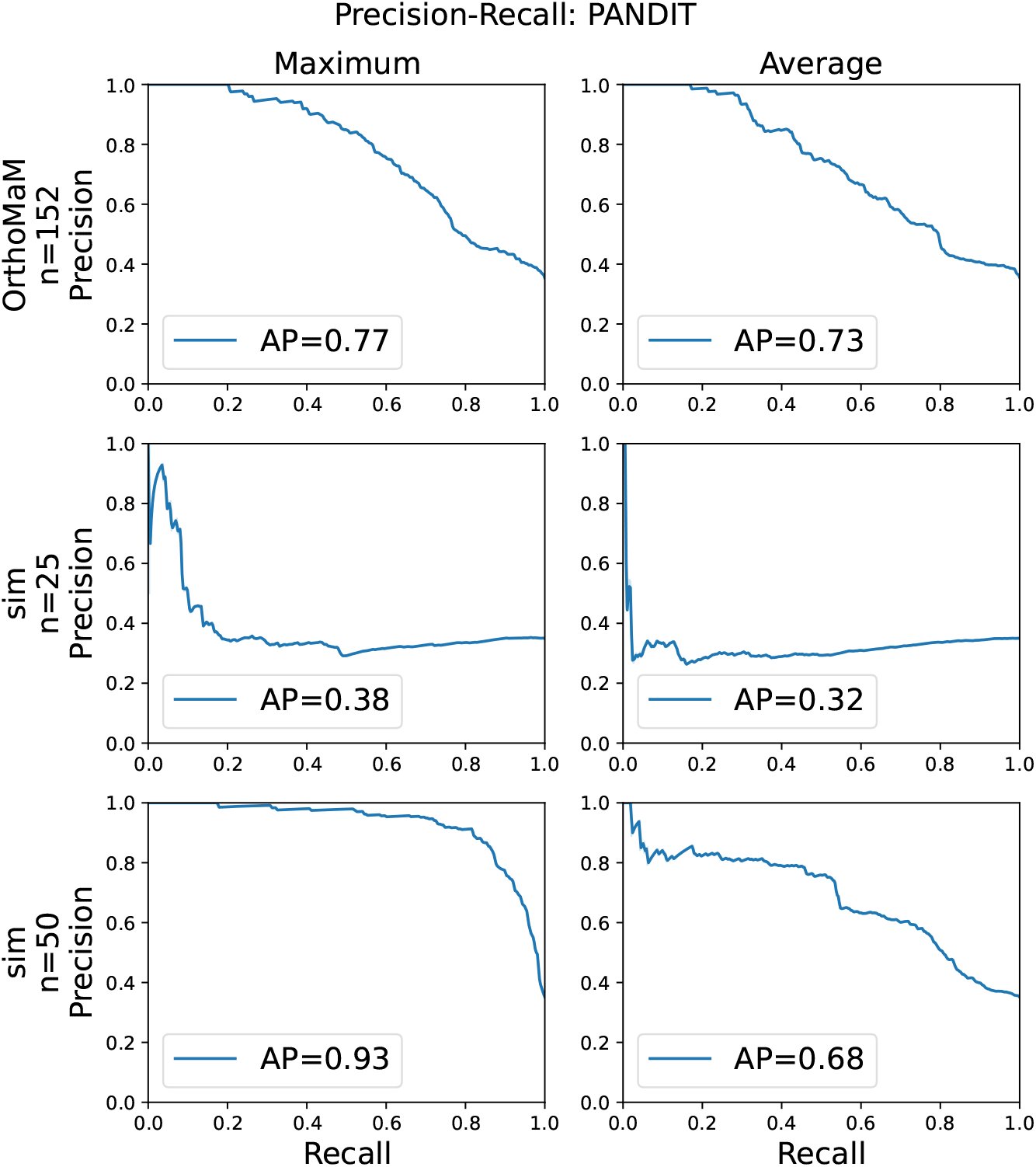
Precision-Recall curve when testing on the PANDIT dataset. Every column represents a different site pooling model (maximum pooling or average pooling), and every row corresponds to a different training data sets: empirical data from the OrthoMaM database in the top row, simulated data with *n* = 25 leaves in the middle row, and simulated data with *n* = 50 leaves in the bottom row. Also shown is the area under the precision-recall curve (AP).

**Figure S2:**
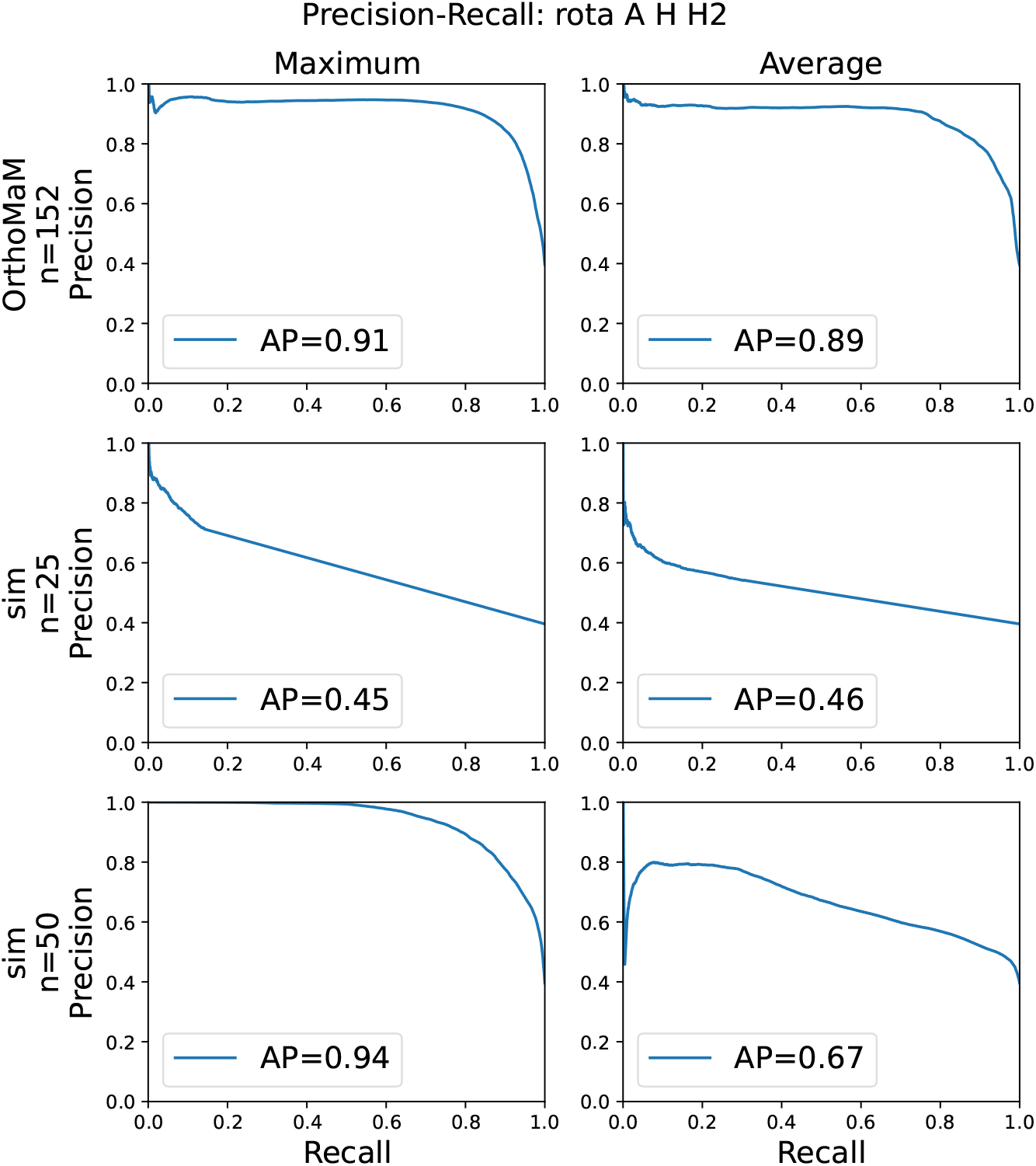
Precision-Recall curve when testing on the rotavirus dataset. Every column represents a different site pooling model (maximum pooling or average pooling), and every row corresponds to a different training data sets: empirical data from the OrthoMaM database in the top row, simulated data with *n* = 25 leaves in the middle row, and simulated data with *n* = 50 leaves in the bottom row. Also shown is the area under the precision-recall curve (AP).

**Figure S3:**
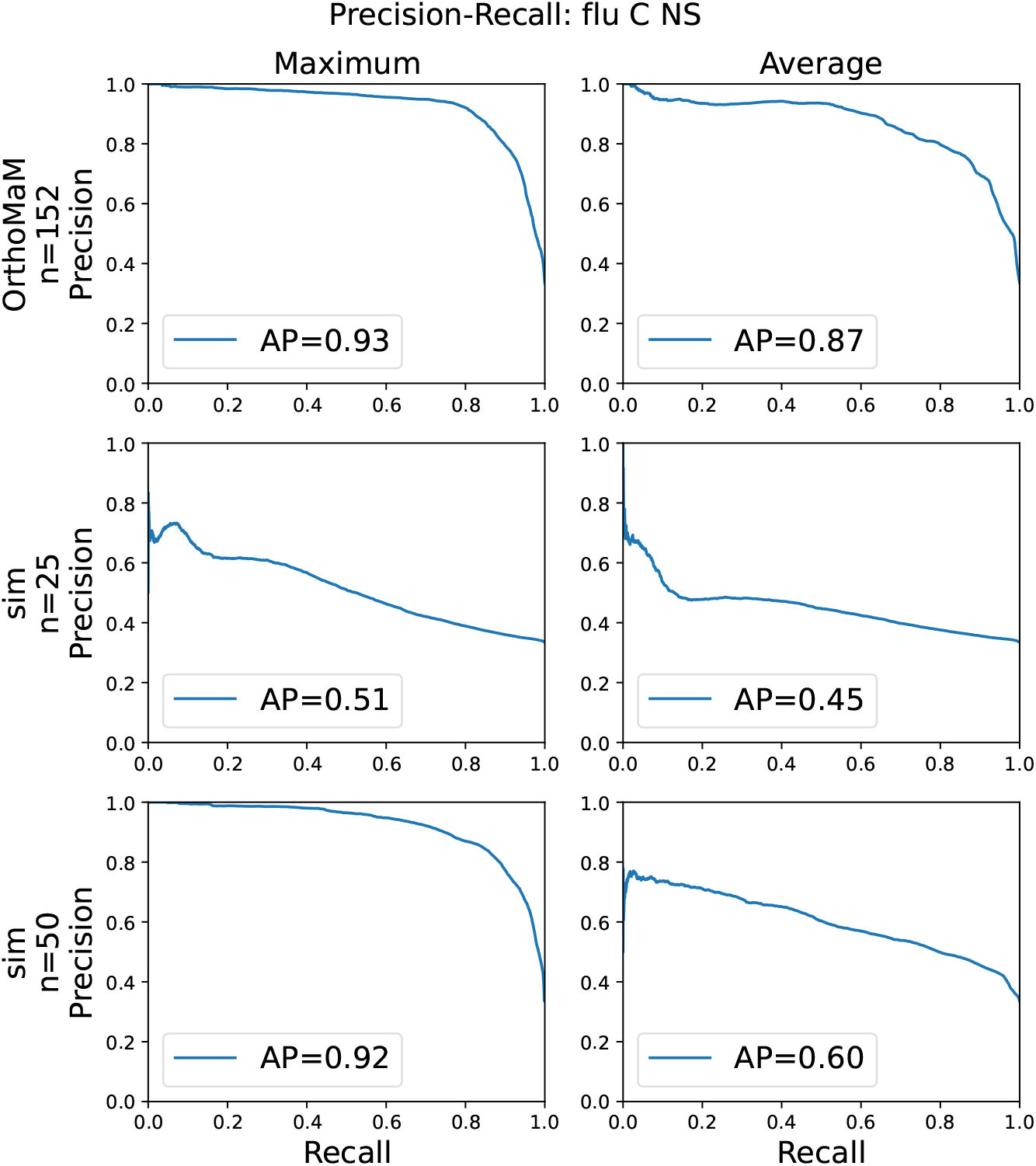
Precision-Recall curve when testing on the influenza C NS dataset. Every column represents a different site pooling model (maximum pooling or average pooling), and every row corresponds to a different training data sets: empirical data from the OrthoMaM database in the top row, simulated data with *n* = 25 leaves in the middle row, and simulated data with *n* = 50 leaves in the bottom row. Also shown is the area under the precision-recall curve (AP).

**Figure S4:**
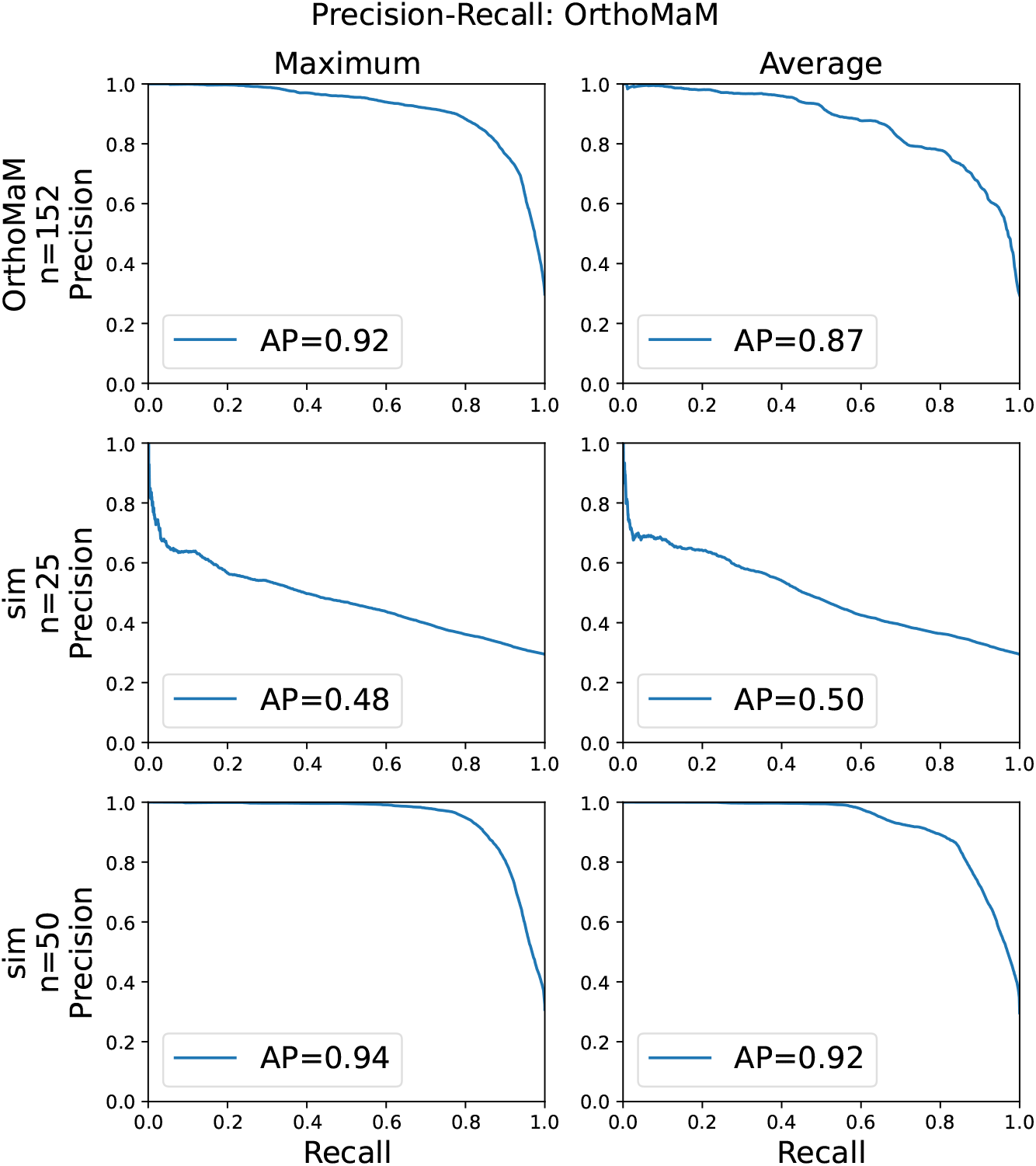
Precision-Recall curve when testing on the OrthoMaM dataset. Every column represents a different site pooling model (maximum pooling or average pooling), and every row corresponds to a different training data sets: empirical data from the OrthoMaM database in the top row, simulated data with *n* = 25 leaves in the middle row, and simulated data with *n* = 50 leaves in the bottom row. Also shown is the area under the precision-recall curve (AP).

**Figure S5:**
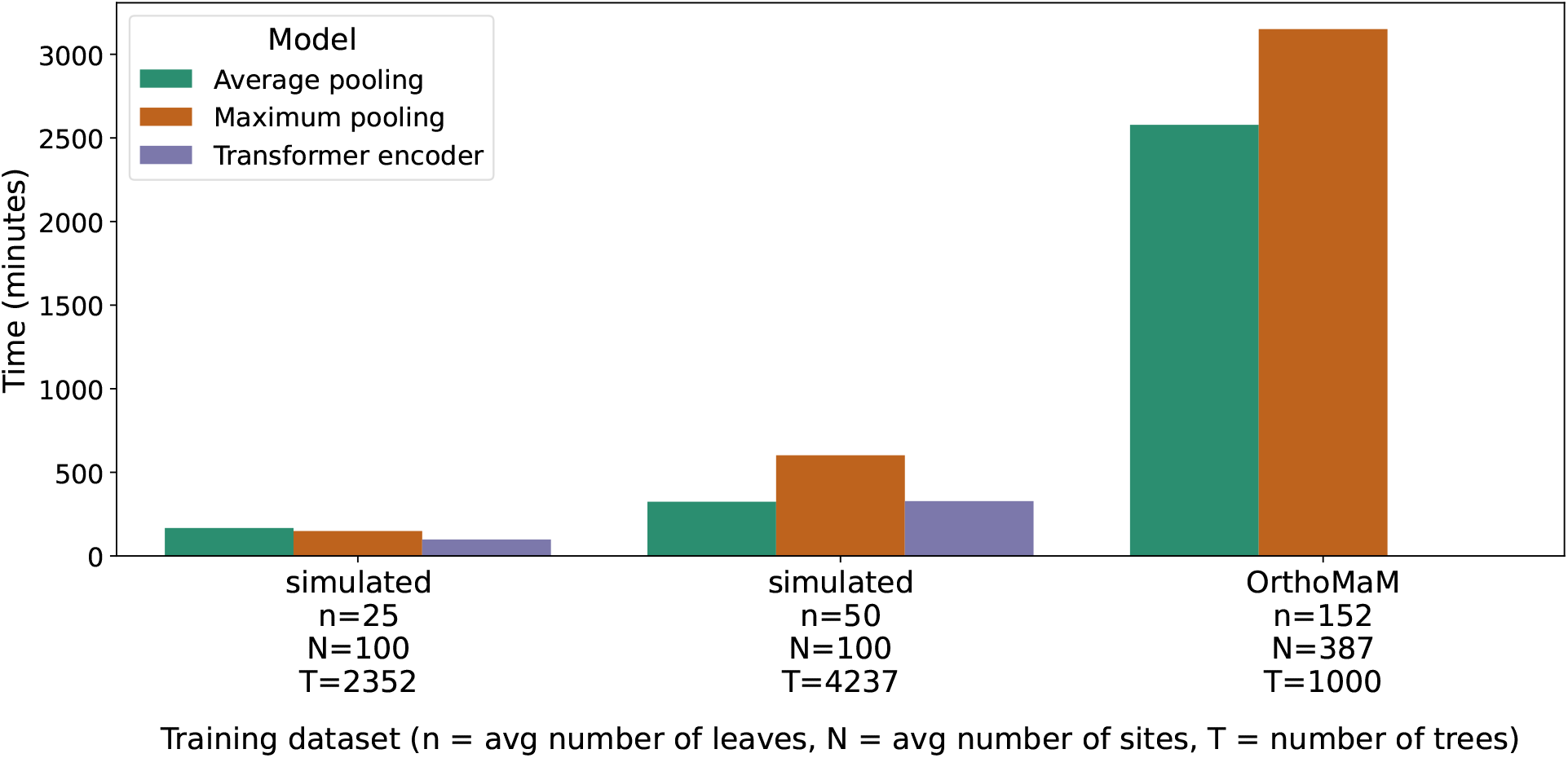
Training wall time in minutes for models trained on simulated data on 25 and 50 leaves and empirical data from the OrthoMaM database. Due to memory restrictions, the OrthoMaM dataset was only used to train the average pooling and maximum pooling model.

**Figure S6:**
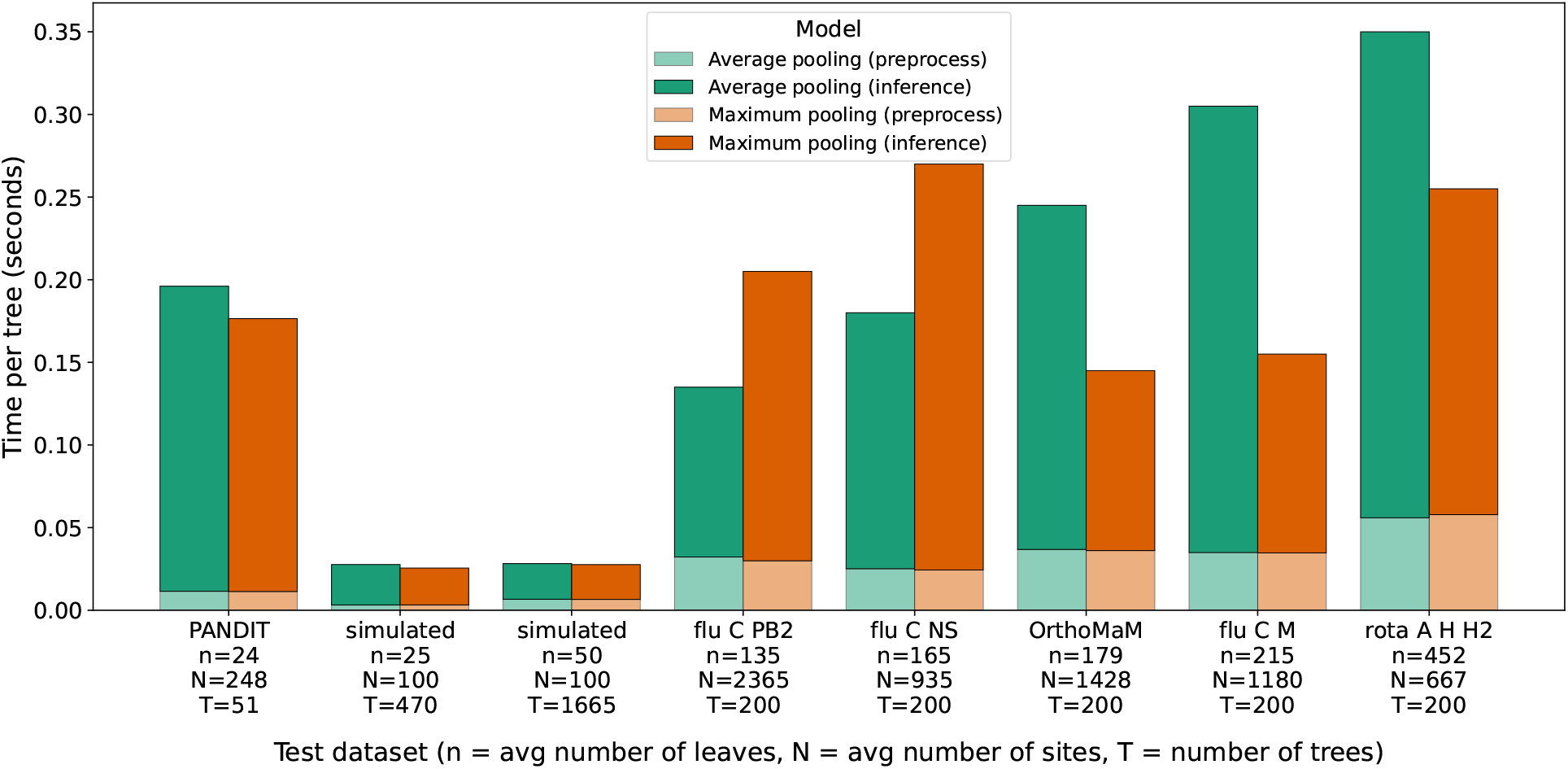
Testing wall time in seconds for our models trained on OrthoMaM training data. Testing datasets are displayed on the x-axis and sorted according to increasing number of leaves in the testing trees. The times include the time required to load the datasets and convert them into the tensor format, labelled as “preprocess”.

**Figure S7:**
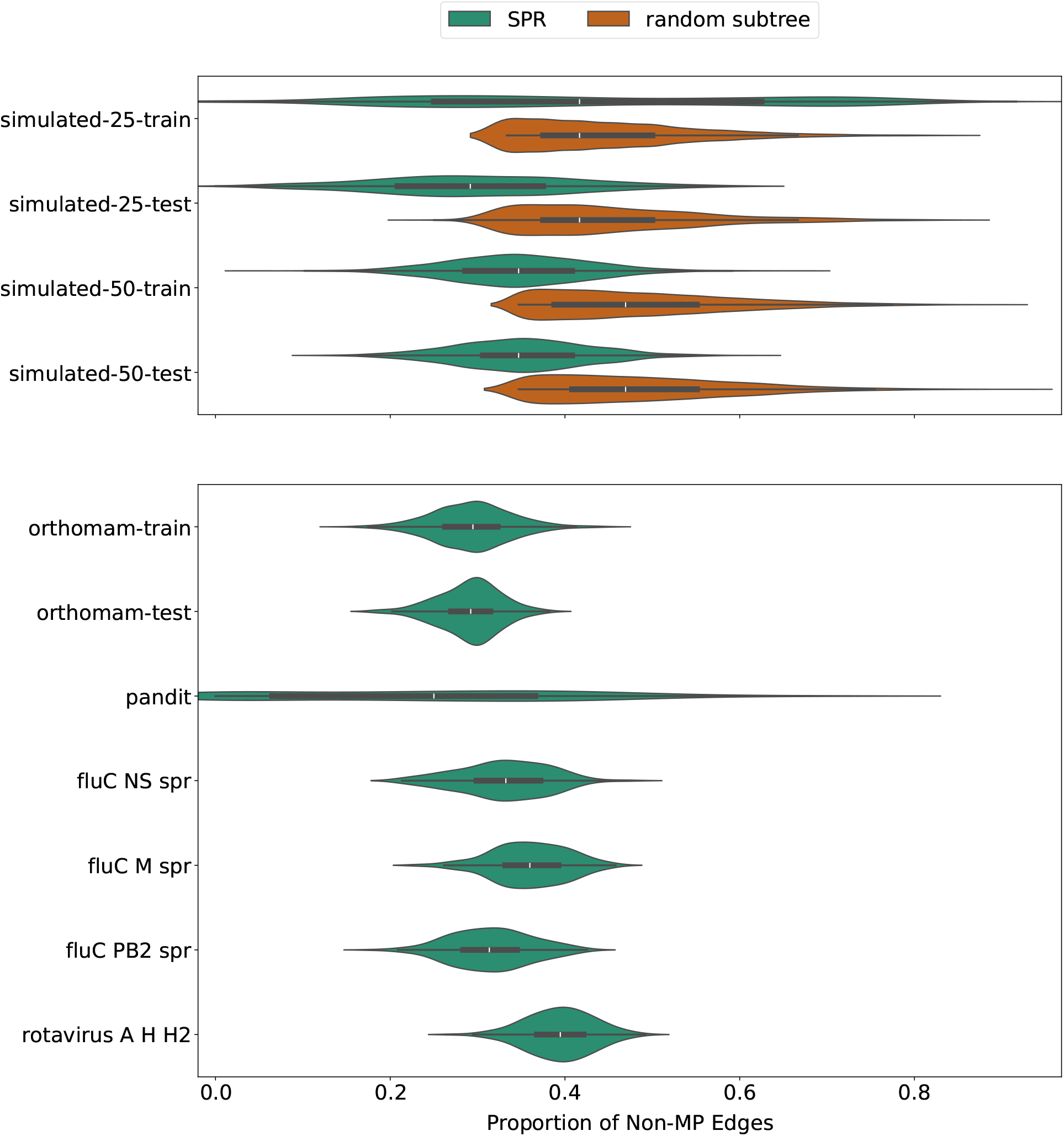
Fraction of non-MP edges in simulated (top) and empirical (bottom) datasets presented in this paper. For simulated datasets we distinguish the different perturbation methods used to introduce non-MP edges (“SPR” vs “random subtree”)

**Figure S8:**
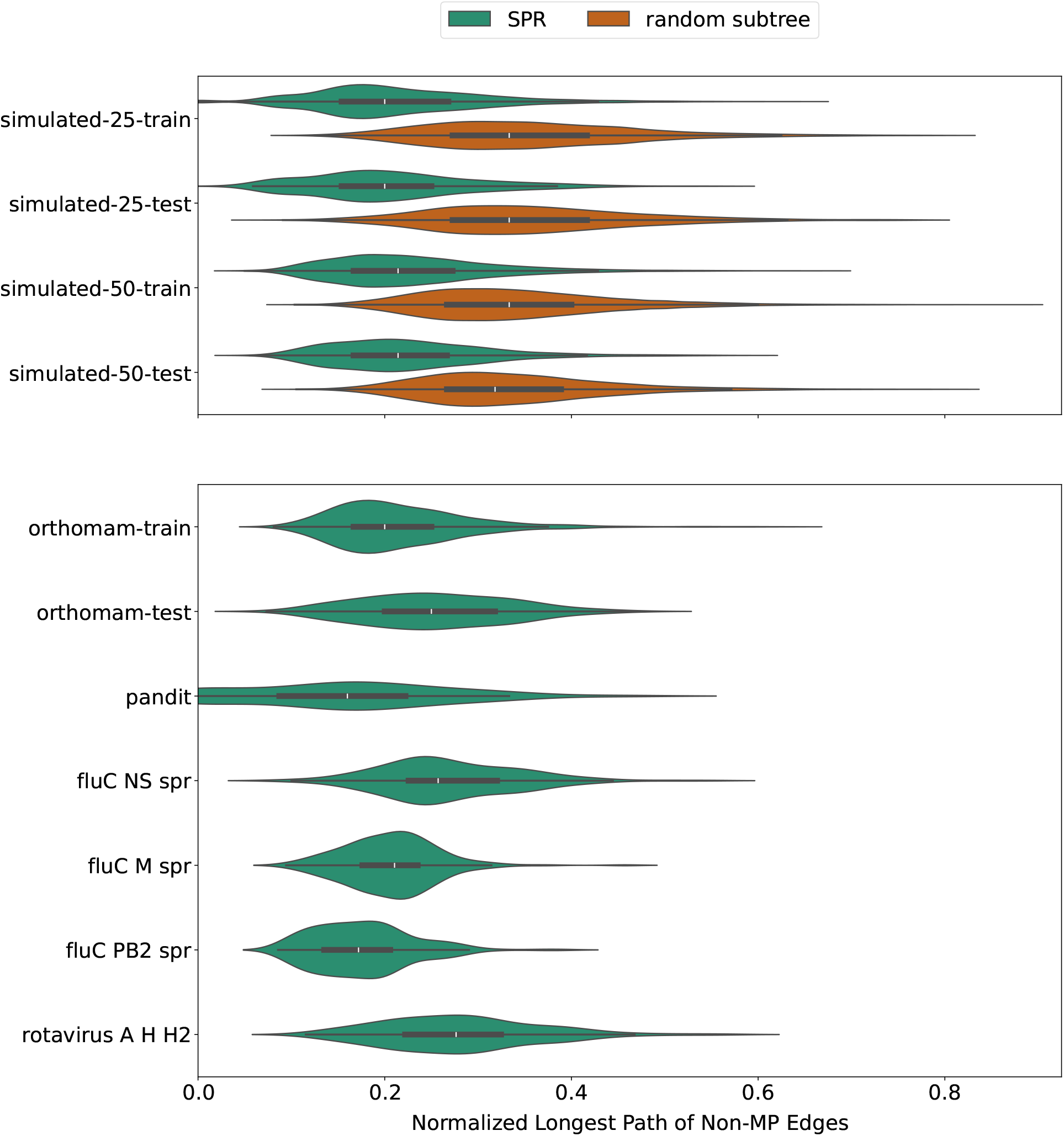
Normalized length of longest non-MP path in simulated (top) and empirical (bottom) datasets presented in this paper. For every tree in the dataset, we take the longest directed path in the tree, i.e. a path is required to be a sub-path of a path from the root to a leaf. For simulated datasets we distinguish the different perturbation methods used to introduce non-MP edges (“SPR” vs “random subtree”)

### Complexity of the MP Edge Problem

**Problem 2** (MP Edge Problem). Let A be an alignment, S_1_|*S*_2_ *a split of A, and k an integer. Is there a tree T for A containing an edge inducing the split S*_1_|*S*_2_ *with* ps*_A_*(*T*) ≤ *k?*

**Theorem 2.** *The* MP Edge Problem *is NP-complete*.

We prove Theorem 2 by reducing the *NP*-complete problem (Day and Sankoff, 1986) of finding a maximum parsimony tree (MP Problem) to the MP Edge Problem.

**Problem S1** (MP Problem). Let A be an alignment and k an integer.

*Is there a tree T for A that has parsimony score* ps*_A_*(*T*) ≤ *k?*

*Proof of Theorem 2.* Given a solution to MP Edge Problem, i.e. a tree *T* for alignment *A* that contains an edge inducing the split *S*_1_|*S*_2_, we can use the Fitch algorithm to compute its parsimony score and verify whether it is less than or equal to the given integer *k*. Therefore, MP Edge Problem is in the class *NP*.

To show *NP*-hardness, we reduce MP Problem to MP Edge Problem. Let *A* be an alignment containing sequences *s*_1_*, s*_2_*, …, s_n_* and *k* be an integer that give us an instance of MP Problem. We use *A^′^* = *A*, the same integer *k*, and the split *S* = {*s*_1_}|{*s*_2_*, s*_3_*, …, s_n_*} as an instance of MP Edge Problem. We now need to prove that there is a tree *T* for *A* with ps*_A_*(*T*) ≤ *k* if and only if there is a tree *T ^′^* for *A^′^* containing an edge inducing the split *S* with ps*_A_′* (*T^′^*) ≤ *k*. With *A^′^* = *A* and *S* being a trivial split that is present in any tree for *A*, it is easy to see that this is true with *T* = *T^′^*.

## Notes

https://doi.org/10.5061/dryad.0zpc867bp

### Competing Interest Statement

The authors have declared no competing interest.

### Summary of Updates

Changed order of Methods and Results and updated results.

